# SunTag-PE: a modular prime editing system enables versatile and efficient genome editing

**DOI:** 10.1101/2023.01.13.524016

**Authors:** Rui Tian, Jiashuo Liu, Ye Chen, Zheying Huang, Lifang Li, Yuyan Wang, Chaoyue Zhong, Tingting Zhao, Zheng Hu

## Abstract

Prime editing (PE) holds tremendous potential in the treatment of genetic diseases because it can install any desired base substitution or local insertion/deletion. However, the full-length PE effector size (6.3-kb) was beyond the packaging capacity of adeno-associated virus (AAV), hindering its clinical transformation. Various splitting strategies have been used to improve its delivery, but always accompanied by compromised PE efficiency. Here, we developed a modular and efficient SunTag-PE system that splits PE effectors into GCN4-nCas9 and single-chain variable fragment (scFv) tethered reverse transcriptase (RT). We observed that SunTag-PEs with 1×GCN4 in the N terminus of nCas9 was the most efficient configuration rather than multiple copies of GCN4. This SunTag-PE strategy achieved editing levels comparable to canonical fused-PE and higher than other split-PE strategies (including sPE and MS2-PE) in both PE2 and PE3 forms with no increase in insertion–deletion (indel) byproducts. Moreover, we successfully validated the modularity of SunTag-PE system in the Cas9 orthologs of SauCas9 and FrCas9. Finally, we employed dual AAVs to deliver SunTag-ePE3 and efficiently corrected the pathogenic mutation in HBB mutant cell line. Collectively, our SunTag-PE system provides an efficient modular splitting strategy for prime editing and further facilitate its transformation in clinics.

## Introduction

Canonical PEs contains two main components, a protein effector consisting of nCas9-H840A tethered with reverse transcriptase (RT)^1,2^ at the C-terminus, and a prime editing guide RNA (pegRNA) containing a primer binding site (PBS) sequence and a RT template^3-5^. It can install desired insertions, deletions, and all 12 possible base-to-base conversions^3,6-8^ without double-strand breaks (DSBs) or donor DNA template, which enables PE to correct the majority of known human genetic disease-related mutations^8-10^.

However, the delivery remains a major obstacle to the clinical translation of PE, as its size exceeds the limited packaging capacity (∼4.8-kb) of AAV^11,12^. To tackle this issue, various splitting strategies have been tested for delivering PEs, including intein-mediated split PEs^13,14^, MS2-PEs^15^, and split-PE^15,16^. But these strategies, while improving delivery efficiency, can also compromise the editing efficiency of PEs.

Here, we developed a modular SunTag-PE strategy that used protein-tagging system^17,18^ to splits the PE effectors into GCN4-nCas9 and scFv-RT, which could be packaged into two separate AAVs and induce efficient editing. We observed that SunTag-PEs could install desired mutations at different endogenous and exogenous loci in HEK293T and HeLa cells, and could achieve higher editing efficiency than canonical fused-PEs and previous different split-PEs. Of note, this SunTag system was also applicable to other orthologous nCas9, which could liberate the restriction of NGG PAM. The modularity of SunTag PE system provides a versatile platform that can advance the effectiveness and broader application scope of PE technology.

## Results

### N-1×GCN4 SunTag-PE was the most efficient configuration

We initially designed SunTag-PE2 by fusing the scFv to the N terminus of Moloney murine leukemia virus reverse transcriptase (M-MLV RT)^19^ and fusing different copies of GCN4 to the N terminus or C terminus of nCas9-H840A^20^. In SunTag-PE system, the scFv-RT was recruited by n×GCN4-nCas9 (the n in n×GCN4 = 1, 2, 3, 5, 10). We designated them as SunTag-PE (n×GCN4-nCas9) and PE-SunTag (nCas9-n×GCN4) based on the domain order (Fig. 1a, b).

**Fig. 1.**
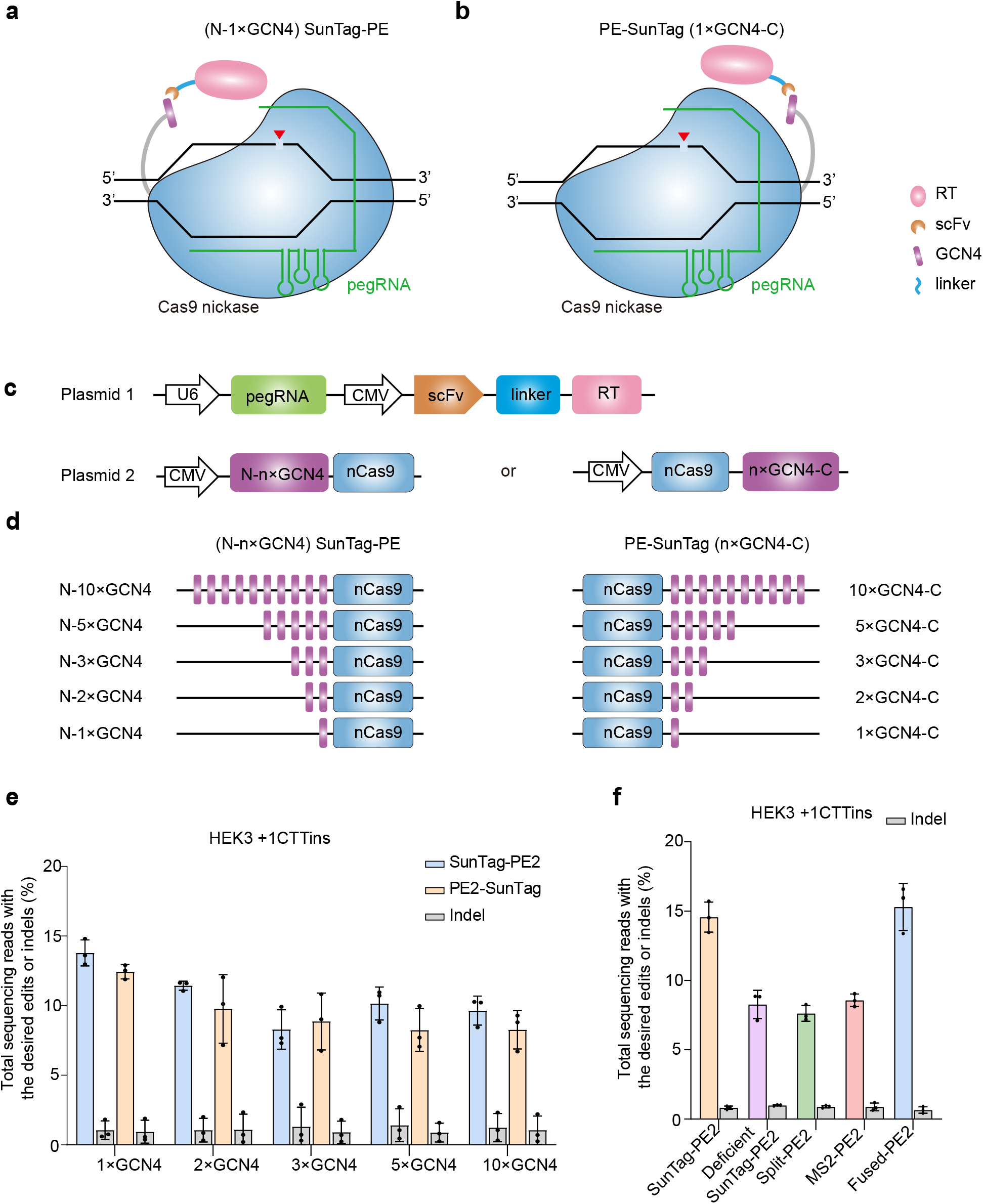
SunTag-PE system enables efficient prime editing in HEK293T cells. **a**, SunTag-PE consists of GCN4 fused to the N-terminus of nCas9. **b**, PE-SunTag consists of GCN4 fused to the C-terminus of nCas9. **c**, Two-plasmid system of SunTag-PE, one plasmid expressing pegRNA and scFv-RT, and the other expressing n×GCN4 tethered nCas9. **d**, Different configurations of SunTag-PE and PE-SunTag (the copy number of GCN4 was 1, 2, 3, 5, and 10). **e**, Comparison of CTT insertion efficiencies of different SunTag-PE2 and PE2-SunTag configurations in HEK3 locus by deep sequencing. **f**, Comparison of CTT insertion efficiencies of SunTag-PE2 with N-1×GCN4, deficient SunTag-PE2, Split-PE2, MS2-PE2, and canonical fused PE2 in HEK3 locus. Data and error bars indicate the mean and s.d. of three independent biological replicates.

Based on the above design principles and reasonable size distribution, we constructed a two-plasmid system, one plasmid expressing pegRNA and scFv-RT (Fig. 1c), and the other expressing n×GCN4 tethered nCas9 (Fig. 1d). To test its feasibility in PE editing, we first co-transfected the above plasmids into HEK293T cells to introduce a 3-nucleotide (nt) CTT insertion in HEK3 endogenous locus. The editing efficiency and indel rate were detected by sanger sequencing (Extended Data Fig. 1) and deep sequencing (Fig. 1e). We observed that the editing efficiency was the highest when there was only one copy of GCN4 tethered to nCas9. And the editing efficiency of GCN4 tethered to the N-terminus of nCas9 was generally better than that of the C-terminus. As shown in Figure 1e and Extended Data Fig. 1, when n×GCN4 was configured at the N-terminal of nCas9, the CTT insertion average rate was 13.8% at 1×GCN4, 11.44% at 2×GCN4, 8.3% at 3×GCN4, 10.16% at 5×GCN4, and 9.64% at 10×GCN4. When n×GCN4 was configured at the C-terminal of nCas9, the CTT insertion average rate was 12.44% at 1×GCN4, 9.77% at 2×GCN4, 8.87% at 3×GCN4, 8.25% at 5×GCN4, and 8.27% at 10×GCN4. The above results were contrary to the previous understandings^17,20^ that the more copies of GCN4 the better efficiency. This phenomenon may be caused by the steric hindrance of neighboring peptide binding sites^17,21^.

### SunTag-PE2 exhibited higher editing efficiency than previous different split PEs

After confirming that the most potent SunTag-tethered configuration was N-1×GCN4-nCas9 (hereinafter refereed as to SunTag-PE), we selected it for further investigation. We first compared the editing efficiency of SunTag-PE with canonical fused-PE^3^, MS2-PE^15^, and split-PE (untethered RT)^15,16^ in the HEK3 locus of HEK293T cells in PE2 form. The data showed that, SunTag-PE2 achieved comparable levels of CTT insertion editing as canonical fused-PE2 (up to 14.58%) and significantly higher than those of split-PE2 (7.63%) and MS2-PE2 (8.57%) with no increase in indel byproducts (Fig. 1f and Extended Data Fig. 2). Furthermore, we confirmed that the predominant editing ability of SunTag-PE2 was derived from the enrichment effect of GCN4-scFv by comparing with the deficient SunTag-PE2 (nCas9 + scFv-RT), which was lack of GCN4 but otherwise identical to SunTag-PE2 (Fig. 1f). Of note, we observed that deficient SunTag-PE2 also achieved comparable editing levels to split-PE2, consistent with previous studies that RT may be able to function as untethered form^15,16^.

### Efficient editing by SunTag-ePE2 at endogenous and exogenous sites

To further improve the editing efficiency of SunTag-PE2, we constructed SunTag-ePE2 by using engineered pegRNAs (epegRNAs) containing a 3′ evoPreQ1 motif^22^. We tested its editing efficiency at four endogenous genomic loci (HEK3, EMX1, FANCF, and RNF2) to install nucleotide transversion, insertion, and deletion in HEK293T cells. We observed 1.5-fold increased efficiency of SunTag-ePE2 compared to SunTag-PE2 in installing CTT insertion in HEK3 locus (14.58% vs. 21.86%, Fig. 1e and 2a). As expected, SunTag-ePE2 could efficiently install desired mutations across all four genomic sites tested with editing level comparable to ePE2 and no increase in indel byproducts (Fig. 2a-d).

**Fig. 2.**
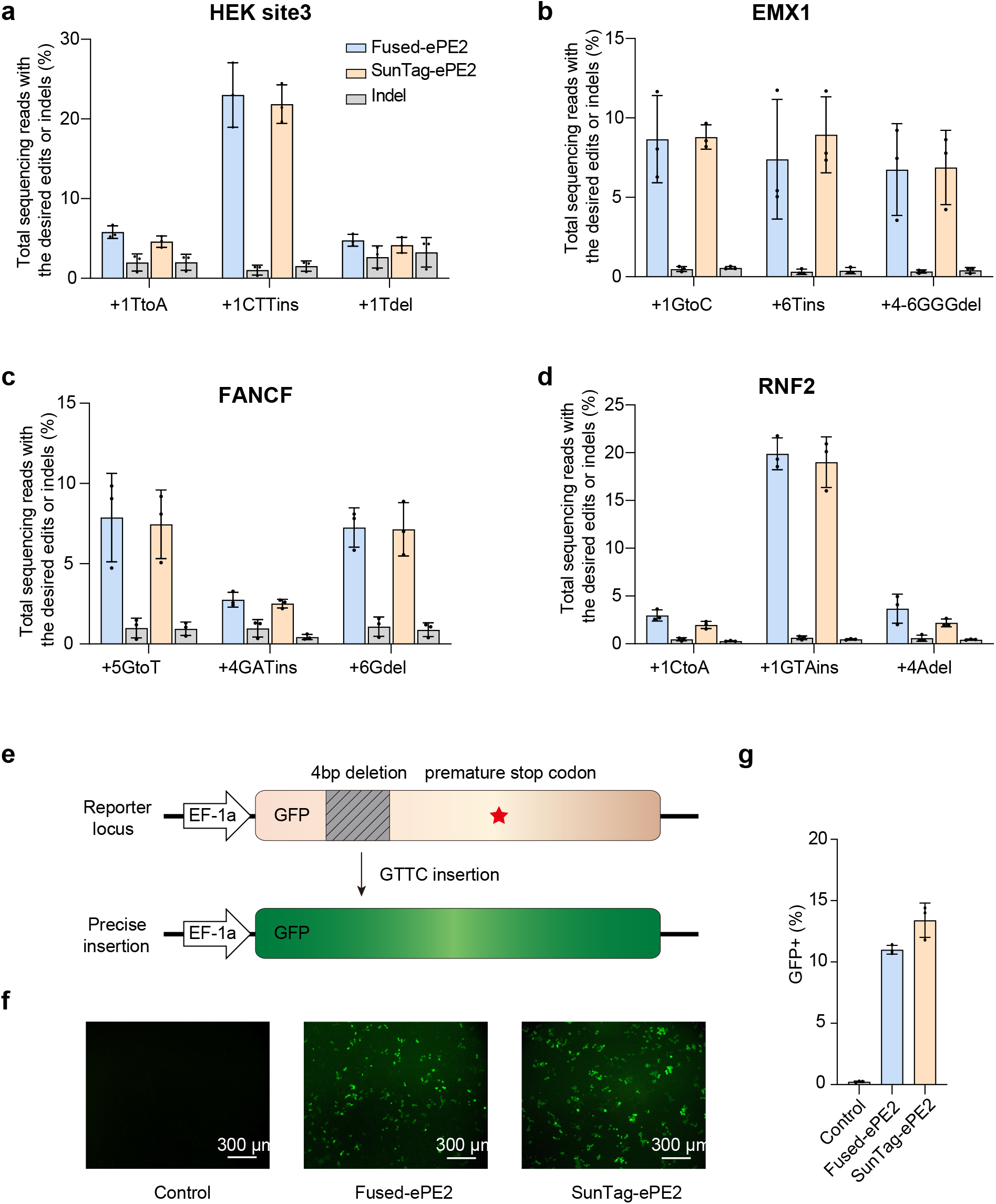
SunTag-ePE2 achieved efficient PE editing comparable to ePE2 at both endogenous and exogenous sites. **a-d**, Comparison of SunTag-ePE2 and ePE2 at four endogenous loci, including HEK3 locus (**a**), EMX1 locus (**b**), FANCF locus (**c**), and RNF2 locus (**d**). Each locus contains three types of prime editing, nucleotide transversion, insertion, and deletion. **e**, The illustration of GFP mutant reporter. It contains a premature stop codon resulting from a 4 nt deletion frameshift mutation in the GFP coding sequence, indicated by red pentagram. Once the GTTC sequence was inserted to the mutation site, it will convert into GFP. **g, f**, The percentage of GFP-positive cells after SunTag-ePE2 and ePE2 editing. Control, reporter cells transfected with PE2 but without pegRNA. Data and error bars indicate the mean and s.d. of three independent biological replicates.

Further, we validated the editing efficiency of SunTag-ePE2 in exogenous site. We first used the piggyBac transposase tool^23^ to establish a green fluorescent protein (GFP)-mutated (GFPm) reporter cell line, containing a premature stop codon resulting from a 4 nt deletion frameshift mutation in the GFP coding sequence^14,24^ (Fig. 2e). Then using SunTag-ePE2 and ePE2 to insert GTTC sequence into the GFP mutation site can convert GFPm into GFP. The rate of GFP positive cells was further calculated by flow cytometer (FCM) to evaluate the PE editing efficiency. The data showed that, the GFP positive cells of SunTag-ePE2 (13.4%) were more than ePE2 (11%), suggesting that the precise editing efficiency of SunTag-ePE2 could be higher than ePE2 (Fig. 2f and Extended Data Fig. 3).

### SunTag-ePE3 achieved comparable levels of editing in HEK293T and higher levels in HeLa cells than ePE3

Previous study showed that PE3 achieve about 3-fold editing efficiency compared with PE2^3^. To demonstrate whether SunTag-PE system was also feasible to PE3, we constructed SunTag-ePE3 system and tested it in four endogenous loci to install desired substitution, insertion, and deletion in both HEK293T and HeLa cells. The data showed that the editing efficiency of SunTag-ePE3 was 2-5-fold higher than that of SunTag-ePE2 across the four genomic loci in HEK293T (Fig. 2 and 3), indicating that the SunTag system was also feasible in PE3 form.

**Fig. 3.**
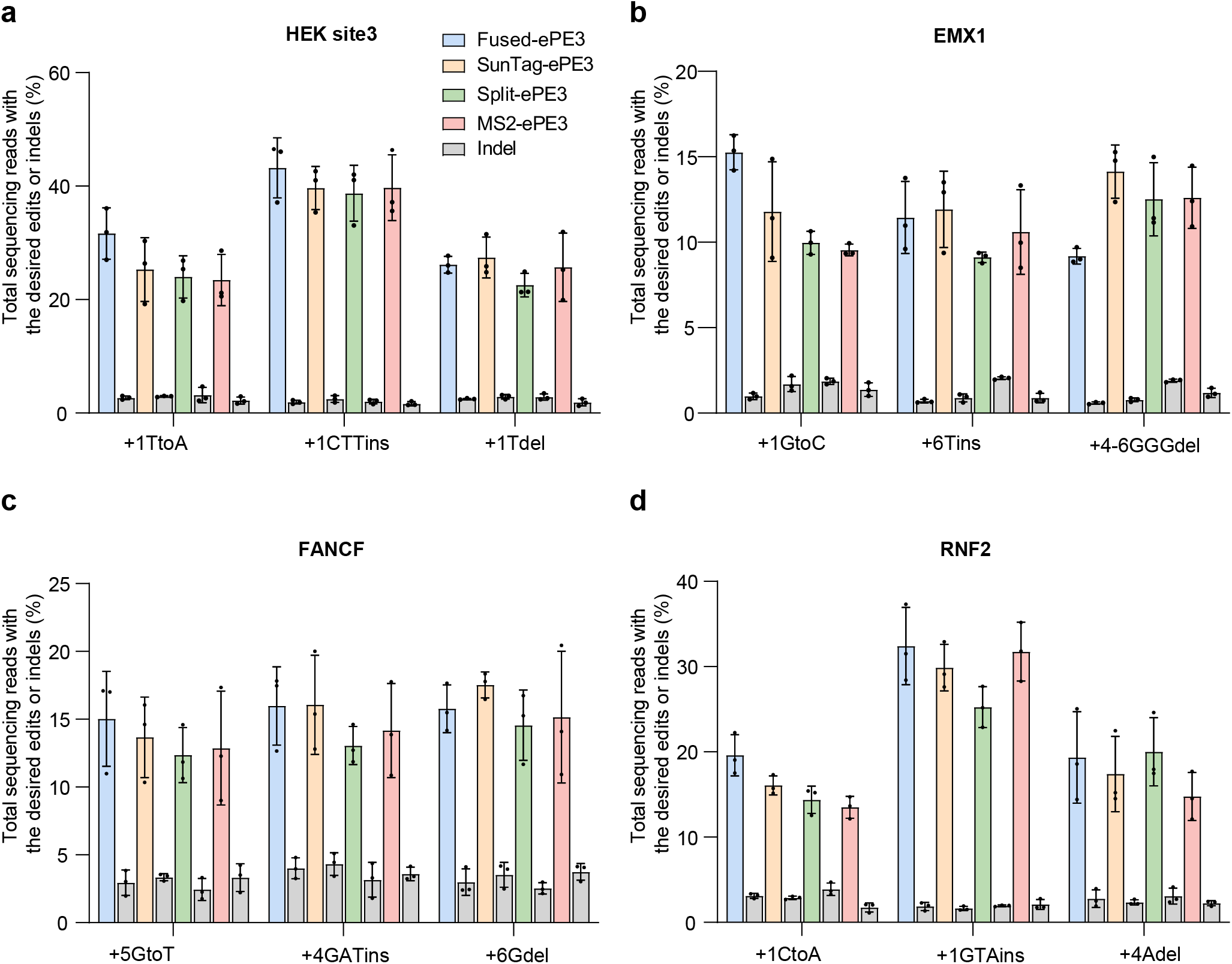
The prime editing efficiency of SunTag-ePE3 was higher than previous split-ePE3 and MS2-ePE3 in HEK293T cells. **a-d**, Comparison of SunTag-ePE3 with canonical fused ePE3, split-ePE3 and MS2-ePE3 at four endogenous loci, including HEK3 locus (**a**), EMX1 locus (**b**), FANCF locus (**c**), and RNF2 locus (**d**) in HEK293T cells. Each locus contains three types of prime editing, nucleotide transversion, insertion, and deletion. Data and error bars indicate the mean and s.d. of three independent biological replicates.

We next compared the SunTag-ePE3 with canonical ePE3, split-ePE3, and MS2-ePE3. As expected, the data in HEK293T cells showed that SunTag-ePE3 achieved comparable editing levels as canonical ePE3 (up to 39.68%) and higher editing levels than split-ePE3 and MS2-ePE3 across all the tested sites in HEK293T cells (Fig. 3).

Since the efficiency of current prime editors varies by cell types^25^, we also tested our SunTag-ePE3 system in HeLa cells at four genomic loci. Surprisingly, we observed that the editing levels of SunTag-ePE3 were higher than canonical ePE3 across all four tested sites for all desired mutations (about 1.1 - 1.8-fold) and with no increase in indel byproducts. (Fig. 4). Specifically, in terms of the substitution efficiency, SunTag-ePE3 was 32.69% in site3 locus, 39.31% in FANCF locus, and 36.68% in RNF2 locus; canonical ePE3 was 29.48% in site3 locus, 33.37% in FANCF locus, and 20.43% in RNF2 locus. In terms of trinucleotide insertion efficiency, SunTag-ePE3 was 51.41% in site3 locus, 55.86% in FANCF locus, and 46.82% in RNF2 locus; canonical ePE3 was 44.16% in site3 locus, 51.05% in FANCF locus, and 31.28% in RNF2 locus. In terms of single nucleotide deletion efficiency, SunTag-ePE3 was 33.27% in site3 locus, 44.99% in FANCF locus, and 26% in RNF2 locus; canonical ePE3 was 26.23% in site3 locus, 40.01% in FANCF locus, and 16.67% in RNF2 locus. The above results indicated that SunTag-PE system was a versatile platform which can accommodate different prime editor versions and exhibited better editing efficiency than canonical fused-PE in specific mammalian cells.

**Fig. 4.**
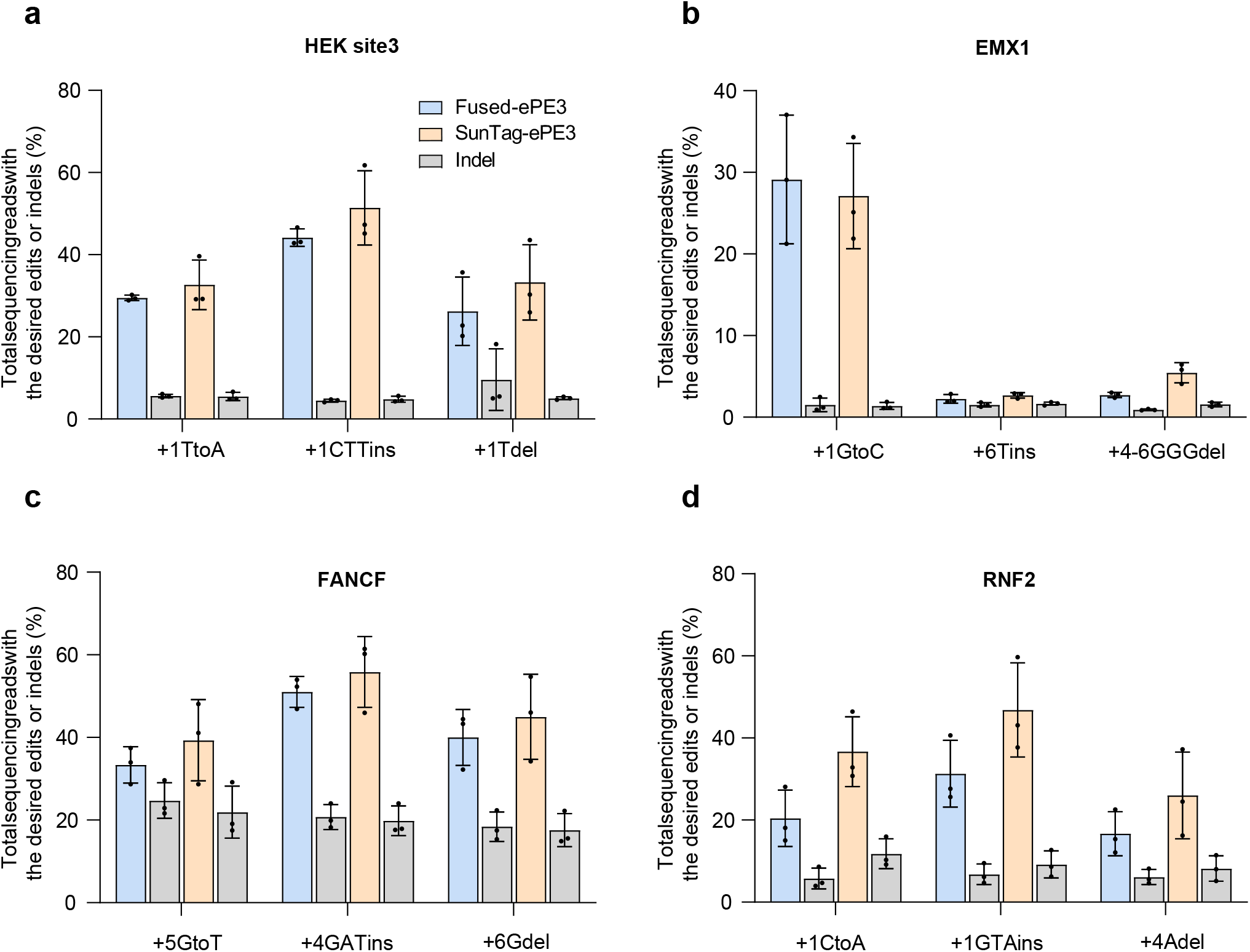
SunTag-ePE3 achieves higher editing level than canonical fused-ePE3 in HeLa cells. **a-d**, Comparison of SunTag-ePE3 with canonical fused ePE3, split-ePE3 and MS2-ePE3 at four endogenous loci, including HEK3 locus (**a**), EMX1 locus (**b**), FANCF locus (**c**), and RNF2 locus (**d**) in HeLa cells. Each locus contains three types of prime editing, nucleotide transversion, insertion, and deletion. Data and error bars indicate the mean and s.d. of three independent biological replicates.

### SunTag-PE system is transferable to other orthologs of Cas9

To further validate the modularity of SunTag-PE system, we constructed SauCas9-SunTag-PE and FrCas9-SunTag-PE to explore whether SunTag-PE was applicable to other orthologs of Cas9. SauCas9 is a smaller Cas9 for all-in-one AAV delivery^26^ and holds tremendous therapeutic potential^27,28^. FrCas9 is a type II-A Cas9 with high cutting efficiency and high fidelity discovered in our laboratory^29^. Notably, we employed active nucleases (SauCas9 and FrCas9) rather than nicking nuclease to construct the DSB-SunTag-PE system. We first examined the prime editing ability of SauCas9-SunTag-PE system in endogenous EMX1 locus and observed efficient editing level in all three types of desired mutations (including transversion, insertion, and deletion) (Extended Data Fig. 5b).

Besides, previous studies revealed that DSB repair will generate undesired consequences, such as pegRNA scaffold integration through nonhomologous end joining (NHEJ) repair pathway^29,30^ (Extended Data Fig. 5a). To improve the precise editing efficiency of DSB-SunTag-PE, we optimized FrCas9-SunTag-PE by removing the RT homology tail of pegRNA and retaining PBS and intended insertions. By using this system, we successfully inserted a short sequence containing a stop codon in the exon 9 of AKT1 gene (Extended Data Fig. 5c). These data suggested that the SunTag-PE system is a modular and easy-to-use platform for constructing PEs with different Cas9 orthologs. Compared with canonical fused-PE, this kind of technology transfer is easier and faster due to modularization, providing more flexibility to prime editing experimental design.

### SunTag-ePE3 delivered by dual-AAV

Having validated the efficient PE editing of SunTag-PE, we further to test whether this system can be delivered with two AAV vectors. First, we used SunTag-PE2 system to install a A•T-to-T•A transversion mutation in HEK293T HBB gene to construct an HBB mutant cell line^31^, mimicking the mutation in sickle cell disease^32^. Then, we encoded the N-1×GCN4-nCas9 (4.67-kb) in one AAV vector and the pegRNAs/ngRNA combination for HBB (T to A) together with scFv-RT in the other (4.36-kb) (Fig.5a). Delivery of both vectors to the HBB mutant cells yielded a mean HBB T-to-A substitution frequency of nearly 25.47 %, whereas delivery of only the pegRNA-ngRNA-scFv-RT did not yield detectable desired editing (Fig. 5C). This modular SunTag-PE address a limitation imposed by size-constrained AAV vectors and avoid additional step of protein reconstitution imposed by intein sequence used previously (split-intein PE2)^8^.

**Fig. 5.**
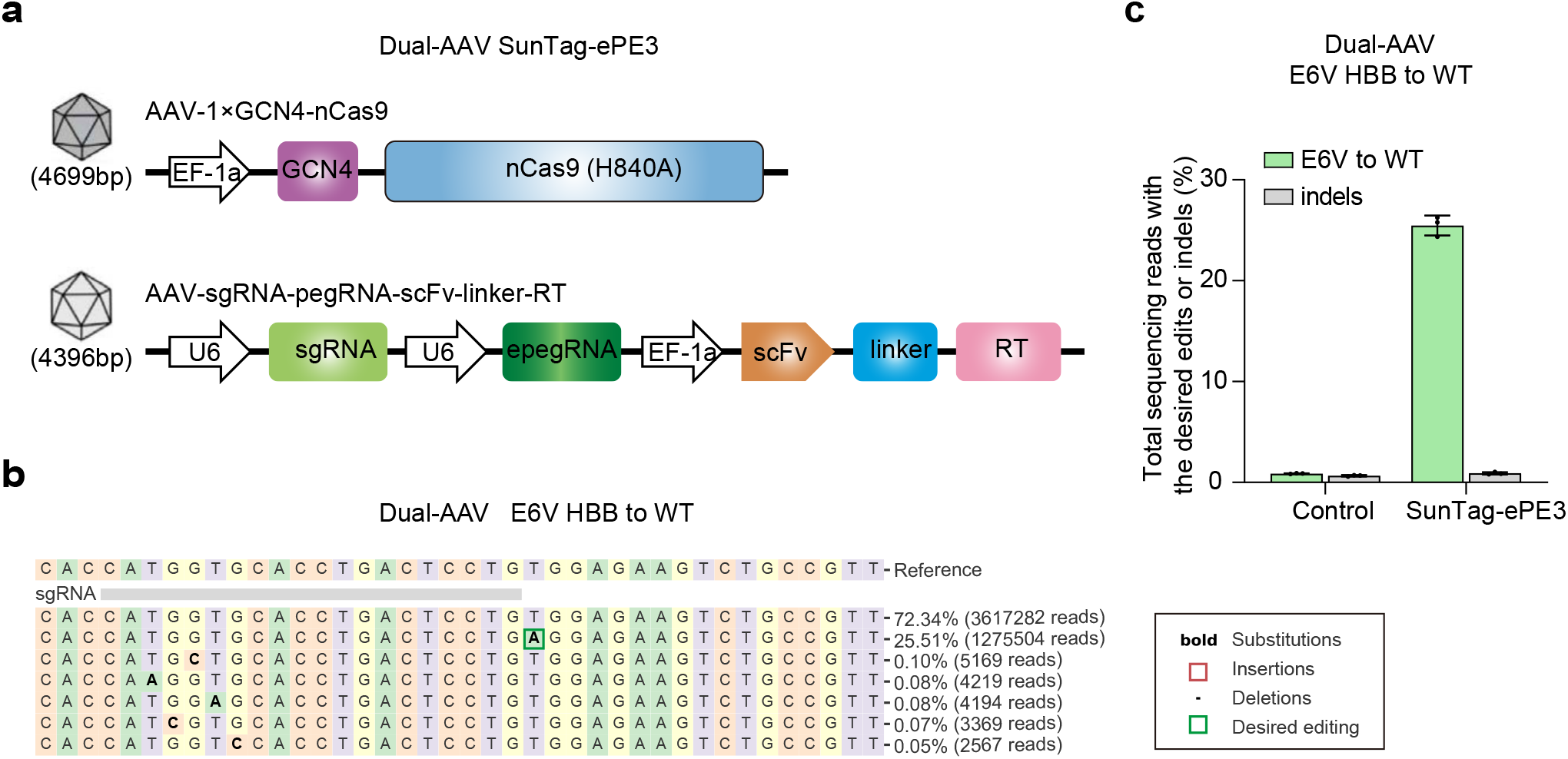
Efficient HBB mutation repair by dual-AAV SunTag-ePE3 in a mutant cell model. **a**, Schematic of SunTag-ePE3 delivered by dual-AAVs. **b**, The HBB mutation repair results of dual-AAV SunTag-ePE3 analyzed by CRISPResso2. **c**, The HBB mutation editing efficiency SunTag-ePE3 in HBB mutant cell model. Data and error bars indicate the mean and s.d. of three independent biological replicates.

## Discussion

In this study, we developed a modular prime editing system, SunTag-PE, which is suitable for different versions of PE, including PE2, ePE2, PE3, and ePE3, and different Cas9 orthologs, such as SauCas9 and FrCas9. During the optimization of the SunTag-PE system, we observed that the 1×GCN4 at the N-terminus of nCas9 has the highest efficiency, which is contrary to the previous report that the more copies of GCN4, the better efficacy (generally 10 GCN4 copies^15,20^ and the highest up to 24 GCN4 copies^17^). We speculated that the decrease in editing efficiency as the copy number of GCN4 increases was caused by the steric clashes of multiple copies of scFv-RT. A recent study also explored the application of SunTag system in prime editing^15^. By fusing a set of 10×GCN4 to the N-terminal or C-terminal of nCas9, the researchers found that the editing efficiency of SunTag-PE with 10×GCN4 was significantly decreased compared to corresponding canonical fused PE3, and concluded that the SunTag system is not suitable for prime editing^15^. However, in this study, we found that SunTag-PE with N-1×GCN4 achieved comparable or even higher editing levels than corresponding canonical fused-PE, and definitely higher than other split-PEs. Therefore, we strongly recommended 1×GCN4 to the N-terminal of nCas9 in the future SunTag-PE research.

We speculated that the high editing efficiency of SunTag-PE system was derived from two aspects. The first was the enrichment effect of GCN4-scFv, which was verified by the better PE editing efficiency of SunTag-PE than that of deficient SunTag-PE (Fig. 1e). The second was the simultaneous function of free soluble RT, as we observed that deficient SunTag-PE can also achieve editing similar to split-PE. Therefore, due to the combination of the two effects, SunTag-PE can obtain editing levels comparable or higher than canonical fused-PE and split-PE.

Notably, the modular and flexible characteristics of SunTag-PE system makes it an easy-to-use and versatile platform for developing new versions of the prime editor. We could explore different versions of RT reverse transcriptase or different pegRNA parameters while keeping GCN4-nCas9 unchanged. This advantage will dramatically facilitate the optimization of PE research. Besides, more and more Cas9 orthologs were validated compatible with SunTag system, such as CjCas9^33^ and SaCas9. In this study, we substituted the nSpCas9 of the SunTag-PE system with active SauCas9 and FrCas9 to construct DSB-SunTag-PE system, and successfully install desired mutation in endogenous loci, not only liberating the SunTag-PE system from the restriction of the NGG PAM^34^, but also providing a new DSB mediated prime editing strategy.

Furthermore, delivery of PEs has been hampered by their large size, SunTag-PE system provides a new splitting strategy to fit in dual AAV vectors^35^. In this study, we employed dual AAVs to deliver SunTag-ePE3 and successfully repaired the pathogenic mutation in an HBB mutant cell line. One AAV vector encoded the N-1×GCN4-nCas9 (4.67-kb) and the other encoded a combination of pegRNAs/ngRNA and scFv-RT (4.36-kb). The smaller size of SunTag-PE leaves more space for the optimization of the length of pegRNA and new type of RT.

In addition, the reduced size of the PE components could improve the yields of mRNA/protein production^36^, easier packaging into the lipid nanoparticles^37,38^, and more efficient entry into cells to achieve higher levels of PE editing. These advantages will further reduce the manufacturing costs and improve the therapeutic efficacy^39^.

Taken together, SunTag-PE system provide a modular splitting PE strategy with high efficiency and flexibility and pave the way for the clinical transformation of effective prime editing.

## Methods

### Plasmids

Plasmids expressing pegRNAs and sgRNA were cloned by Gibson assembly. sgRNA oligos (including 20-bp U6 promoter, spacer, and 20-bp scaffold) were synthesized from Genewiz (Suzhou) and cloned into the linearized PXZ vector by Gibson assembly using Gibson Assembly Master Mix (New England Biolabs). pegRNA plasmids were constructed by using the plasmid (Addgene #132778) as PCR template with the forward primers (containing spacers) and reverse primers (containing PBS and RTT sequences). Then, the PCR product was subsequently cloned into the linearized PXZ vector by Gibson assembly. epegRNA linker sequences were designed via pegLIT^22^ to minimize base pairing between linker and the remainder of the pegRNA. The sgRNA scaffold fragments with MS2 sequences were synthesized and followed by Gibson assembly. Sequences of pegRNAs and nicking sgRNAs were listed in Supplementary Table 1.

PE2 plasmid was obtained from Addgene (#132775). The scFv sequence was derived from plasmid (Addgene #60904) via PCR. The scFv-RT fragment was constructed by replacing nCas9 with the scFv sequence in PE2 plasmid, and this intermediate vector was subsequently assembled with PCR amplified U6-pegRNA fragment, finally obtained U6-pegRNA-CMV-scFv-RT plasmids. n×GCN4 sequences were derived from plasmid (Addgene #113022) via PCR, then N terminus or C terminus n×GCN4 of nCas9 plasmids were constructed, respectively. The RT plasmid was constructed by amplifying nCas9 fragment from PE2 via PCR. The MCP sequence was derived from plasmid (Addgene #85416) via PCR. The MCP-RT plasmid was prepared by replaced nCas9 with MCP to the N-terminal of M-MLV RT. The SaCas9 sequence was derived from Addgene #124844 via PCR. FrCas9 sequence was synthesized by Genewiz. All plasmids were verified by Sanger sequencing and purified with kit (QIAGEN) according to the manufacture’ protocol.

### Cell culture and transfection

Human Embryonic Kidney 293T (HEK293T) cell line and HeLa cervical cancer cell line were purchased from ATCC. These cells were cultured in Dulbecco’s Modified Eagle Medium (DMEM) supplemented with 10%FBS (Gibco) and incubated at 37 °C with 5% CO2. HEK293T cells were transfected using X-tremeGENE™ HP DNA Transfection Reagent (Roche, 6366244001) according to the manufacturer’s protocol. 1×10^5^ HEK293T cells were seeded per well in a 12-well plate overnight. For canonical fused-PE experiment, 330 ng of pegRNA, 110 ng of nicking sgRNA and 1 μg of PE2 plasmids were used for each well. The same molar amounts of plasmids were used for SunTag-PE, deficient SunTag-PE, Split-PE, and MS2-PE. 2×10^5^ HeLa cells were electroporated using a Lonza 4-D Nucleofector with the SE Kit and the CN-114 program. All of the transfected cells were further cultured for 72 h and then were harvested for genomic DNA extraction.

### AAV vector production and transduction

AAV vectors (AAV DJ capsids) were packaged and produced as previouslydescribed^40^. Vector titers were determined by qPCR with ITR-specific primers. For AAV transduction, 5×10^4^ cells were seeded in 24-well plates and infected with dual AAVs at 10^6^ MOI. Five days post-transduction, cells were harvested for genomic DNA extraction.

### Genomic DNA extraction and PCR

Genomic DNA was extracted with EasyPure® Genomic DNA Kit (Transgen, TEE101-01) according to the manufacturer’s instruction. Then 200 ng of genomic DNA was used in a PCR reaction using 2×EasyTaq® PCR SuperMix (Transgen, AS111-01) and primers surrounding the targeting region. The primer sequences used for amplification of edited locus were list in Supplementary Table 2.

### Sanger sequencing and analysis using EditR

The PCR products were determined by gel electrophoresis and sequenced by Sanger sequencing (Sangon, Guangzhou). The results were quantified using the online EditR tool (http://baseeditr.com)^41^.

### Deep sequencing and data analysis

The amplicon PCR products for next-generation sequencing were obtained through two step PCR reaction. In brief, 200 ng genomic DNA was used as a template for PCR amplification with KAPA HiFi HotStart ReadyMix (KAPA Biosystems, KK2602) and primer pairs containing an internal locus-specific region and an outer Illumina-compatible adaptor sequence. A second PCR reaction targeting the outer-adaptor sequence was performed to append unique dual index to each amplicon. Amplification was performed with 25 cycles for the first PCR and 11 cycles for the second PCR. The sequences of the primers were listed in Supplementary Table 3. Then, the libraries were sequenced in a MGI2000 platform with paired-end 150 after library adaptor transformation. For data analysis, after quality control of the raw sequencing data by fastp with default parameters^42^, pair-end reads were merged into one read by FLASH software^43^. Then, prime editing efficiency was determined by aligning amplicon reads to a reference sequence using CRISPResso2^44^. The fraction of precise editing was calculated as the desired editing reads /total sequencing reads. The results were loaded into GraphPad Prism 8.4 for data visualization.

### Flow cytometry analysis

Flow cytometry analysis was performed on day 3 after transfection. Cells were washed with PBS and digested by 0.25% trypsin, followed by centrifugation and resuspension in PBS with 2% FBS. The GFP signal was detected by FITC channel using CytoFLEX Flow Cytometer (Beckman). 10,000 events were counted from each sample for *FCM* analysis. Data were analyzed by FlowJo v10 software. *FCM* gating examples for reporter cells were shown in Extended Data Fig. 3.

## Data availability

Plasmids encoding constructs used in this study will be deposited at Addgene. All targeted amplicon sequencing data will be deposited at the National Center of Biotechnology Information’s Sequence Read Archive.

## Acknowledgements

This work was supported by The National Natural Science Foundation of China (Grant no. 32171465 and 82102392); General Program of Natural Science Foundation of Guangdong Province of China (Grant no. 2021A1515012438); Guangdong Basic and Applied Basic Research Foundation (Grant no. 2020A1515110170); the National Ten Thousand Plan-Young Top Talents of China (Grant no. 80000-41180002).

## Author contributions

R.T. and J.L. contributed equally and designed the research, performed experiments and analyzed data. Y.C., Zhe.H., L.L., Chao.Z., T.Z., and Chu.Z performed experiments. Y.W. analyzed the NGS data. Z.H. designed and supervised the research. R.T., J.L., and Z.H. wrote the paper.

## Additional information

Correspondence and requests for materials should be addressed to R.T. or Z.H.

## References

1 Levin, H. L. It’s prime time for reverse transcriptase. Cell 88, 5–8, doi:10.1016/s0092-8674(00)81851-6 (1997).

2 Huber, L. B., Betz, K. & Marx, A. Reverse Transcriptases: From Discovery and Applications to Xenobiology. Chembiochem, e202200521, doi:10.1002/cbic.202200521 (2022).

3 Anzalone, A. V. et al. Search-and-replace genome editing without double-strand breaks or donor DNA. Nature 576, 149–157, doi:10.1038/s41586-019-1711-4 (2019).

4 Sheridan, C. Gene editing enters ‘prime’ time. Nat Biotechnol, doi:10.1038/d41587-019-00032-5 (2019).

5 Yang, L., Yang, B. & Chen, J. One Prime for All Editing. Cell 179, 1448–1450, doi:10.1016/j.cell.2019.11.030 (2019).

6 Chemello, F. et al. Precise correction of Duchenne muscular dystrophy exon deletion mutations by base and prime editing. Sci Adv 7, doi:10.1126/sciadv.abg4910 (2021).

7 Lin, Q. et al. Prime genome editing in rice and wheat. Nat Biotechnol 38, 582–585, doi:10.1038/s41587-020-0455-x (2020).

8 Liu, P. et al. Improved prime editors enable pathogenic allele correction and cancer modelling in adult mice. Nat Commun 12, 2121, doi:10.1038/s41467-021-22295-w (2021).

9 Chen, P. J. et al. Enhanced prime editing systems by manipulating cellular determinants of editing outcomes. Cell 184, 5635-5652.e5629, doi:10.1016/j.cell.2021.09.018 (2021).

10 Hampton, T. DNA Prime Editing: A New CRISPR-Based Method to Correct Most Disease-Causing Mutations. JAMA 323, 405–406, doi:10.1001/jama.2019.21827 (2020).

11 Wu, Z., Yang, H. & Colosi, P. Effect of genome size on AAV vector packaging. Mol Ther 18, 80–86, doi:10.1038/mt.2009.255 (2010).

12 Grieger, J. C. & Samulski, R. J. Packaging capacity of adeno-associated virus serotypes: impact of larger genomes on infectivity and postentry steps. J Virol 79, 9933–9944, doi:10.1128/JVI.79.15.9933-9944.2005 (2005).

13 Zheng, C. et al. A flexible split prime editor using truncated reverse transcriptase improves dual-AAV delivery in mouse liver. Mol Ther 30, 1343–1351, doi:10.1016/j.ymthe.2022.01.005 (2022).

14 Zhi, S. et al. Dual-AAV delivering split prime editor system for in vivo genome editing. Mol Ther 30, 283–294, doi:10.1016/j.ymthe.2021.07.011 (2022).

15 Liu, B. et al. A split prime editor with untethered reverse transcriptase and circular RNA template. Nat Biotechnol, doi:10.1038/s41587-022-01255-9 (2022).

16 Grunewald, J. et al. Engineered CRISPR prime editors with compact, untethered reverse transcriptases. Nat Biotechnol, doi:10.1038/s41587-022-01473-1 (2022).

17 Tanenbaum, M. E., Gilbert, L. A., Qi, L. S., Weissman, J. S. & Vale, R. D. A protein-tagging system for signal amplification in gene expression and fluorescence imaging. Cell 159, 635–646, doi:10.1016/j.cell.2014.09.039 (2014).

18 Ye, H., Rong, Z. & Lin, Y. Live cell imaging of genomic loci using dCas9-SunTag system and a bright fluorescent protein. Protein Cell 8, 853–855, doi:10.1007/s13238-017-0460-0 (2017).

19 Shinnick, T. M., Lerner, R. A. & Sutcliffe, J. G. Nucleotide sequence of Moloney murine leukaemia virus. Nature 293, 543–548, doi:10.1038/293543a0 (1981).

20 Jiang, W. et al. BE-PLUS: a new base editing tool with broadened editing window and enhanced fidelity. Cell Res 28, 855–861, doi:10.1038/s41422-018-0052-4 (2018).

21 Orozco, C. T. et al. Mechanistic insights into the rational design of masked antibodies. MAbs 14, 2095701, doi:10.1080/19420862.2022.2095701 (2022).

22 Nelson, J. W. et al. Engineered pegRNAs improve prime editing efficiency. Nat Biotechnol 40, 402–410, doi:10.1038/s41587-021-01039-7 (2022).

23 Li, X. et al. piggyBac transposase tools for genome engineering. Proc Natl Acad Sci U S A 110, E2279–2287, doi:10.1073/pnas.1305987110 (2013).

24 Paillusson, A., Hirschi, N., Vallan, C., Azzalin, C. M. & Muhlemann, O. A GFP-based reporter system to monitor nonsense-mediated mRNA decay. Nucleic Acids Res 33, e54, doi:10.1093/nar/gni052 (2005).

25 Chen, P. J. et al. Enhanced prime editing systems by manipulating cellular determinants of editing outcomes. Cell 184, 5635–5652 e5629, doi:10.1016/j.cell.2021.09.018 (2021).

26 Ibraheim, R. et al. All-in-one adeno-associated virus delivery and genome editing by Neisseria meningitidis Cas9 in vivo. Genome Biol 19, 137, doi:10.1186/s13059-018-1515-0 (2018).

27 Friedland, A. E. et al. Characterization of Staphylococcus aureus Cas9: a smaller Cas9 for all-in-one adeno-associated virus delivery and paired nickase applications. Genome Biol 16, 257, doi:10.1186/s13059-015-0817-8 (2015).

28 Ibraheim, R. et al. Self-inactivating, all-in-one AAV vectors for precision Cas9 genome editing via homology-directed repair in vivo. Nat Commun 12, 6267, doi:10.1038/s41467-021-26518-y (2021).

29 Tao, R. et al. WT-PE: Prime editing with nuclease wild-type Cas9 enables versatile large-scale genome editing. Signal Transduct Target Ther 7, 108, doi:10.1038/s41392-022-00936-w (2022).

30 Adikusuma, F. et al. Optimized nickase- and nuclease-based prime editing in human and mouse cells. Nucleic Acids Res 49, 10785–10795, doi:10.1093/nar/gkab792 (2021).

31 Shih, H. C. et al. Rapid identification of HBB gene mutations by high-resolution melting analysis. Clin Biochem 42, 1667–1676, doi:10.1016/j.clinbiochem.2009.07.017 (2009).

32 Park, S. H. et al. Highly efficient editing of the beta-globin gene in patient-derived hematopoietic stem and progenitor cells to treat sickle cell disease. Nucleic Acids Res 47, 7955–7972, doi:10.1093/nar/gkz475 (2019).

33 Zhang, X. et al. MiniCAFE, a CRISPR/Cas9-based compact and potent transcriptional activator, elicits gene expression in vivo. Nucleic Acids Res 49, 4171–4185, doi:10.1093/nar/gkab174 (2021).

34 Karvelis, T. et al. Rapid characterization of CRISPR-Cas9 protospacer adjacent motif sequence elements. Genome Biol 16, 253, doi:10.1186/s13059-015-0818-7 (2015).

35 Bulcha, J. T., Wang, Y., Ma, H., Tai, P. W. L. & Gao, G. Viral vector platforms within the gene therapy landscape. Signal Transduct Target Ther 6, 53, doi:10.1038/s41392-021-00487-6 (2021).

36 Rohner, E., Yang, R., Foo, K. S., Goedel, A. & Chien, K. R. Unlocking the promise of mRNA therapeutics. Nat Biotechnol 40, 1586–1600, doi:10.1038/s41587-022-01491-z (2022).

37 Hou, X., Zaks, T., Langer, R. & Dong, Y. Lipid nanoparticles for mRNA delivery. Nat Rev Mater 6, 1078–1094, doi:10.1038/s41578-021-00358-0 (2021).

38 Cullis, P. R. & Hope, M. J. Lipid Nanoparticle Systems for Enabling Gene Therapies. Mol Ther 25, 1467–1475, doi:10.1016/j.ymthe.2017.03.013 (2017).

39 Demirci, S. et al. Advances in CRISPR Delivery Methods: Perspectives and Challenges. CRISPR J 5, 660–676, doi:10.1089/crispr.2022.0051 (2022).

40 Gao, Z. et al. A truncated reverse transcriptase enhances prime editing by split AAV vectors. Mol Ther, doi:10.1016/j.ymthe.2022.07.001 (2022).

41 Kluesner, M. G. et al. EditR: A Method to Quantify Base Editing from Sanger Sequencing. Crispr j 1, 239–250, doi:10.1089/crispr.2018.0014 (2018).

42 Chen, S., Zhou, Y., Chen, Y. & Gu, J. fastp: an ultra-fast all-in-one FASTQ preprocessor. Bioinformatics 34, i884–i890, doi:10.1093/bioinformatics/bty560 (2018).

43 Magoc, T. & Salzberg, S. L. FLASH: fast length adjustment of short reads to improve genome assemblies. Bioinformatics 27, 2957–2963, doi:10.1093/bioinformatics/btr507 (2011).

44 Clement, K. et al. CRISPResso2 provides accurate and rapid genome editing sequence analysis. Nat Biotechnol 37, 224–226, doi:10.1038/s41587-019-0032-3 (2019).

