## Supplementary Information 1 for "SunTag-PE: a modular prime editing system enables versatile and efficient genome editing"

**Supplementary Figures and Figure legends**

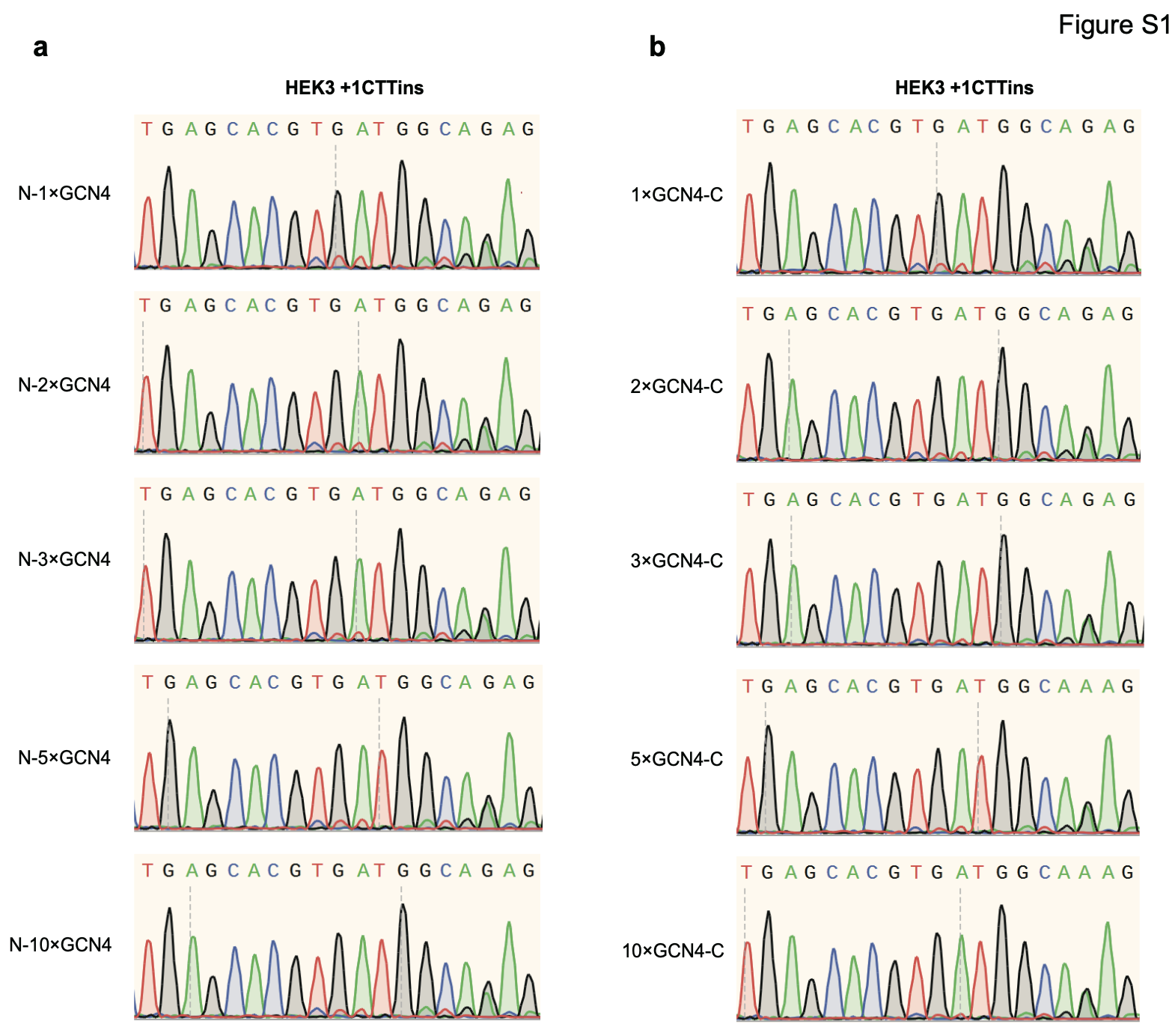

**Fig S1. Sanger sequencing results of SunTag-PE2 installing HEK3 +1CTT insertion in HEK293T cells in different configurations**. **a**. The sanger sequencing results of CTT insertion of SunTag-PE2 with N-n×GCN4 (n = 1, 2, 3, 5, and 10) tethered to the N terminus of nCas9-H840A. **b**. The sanger sequencing results of CTT insertion of SunTag-PE2 with C-n×GCN4 (n = 1, 2, 3, 5, and 10) tethered to the C terminus of nCas9-H840A.

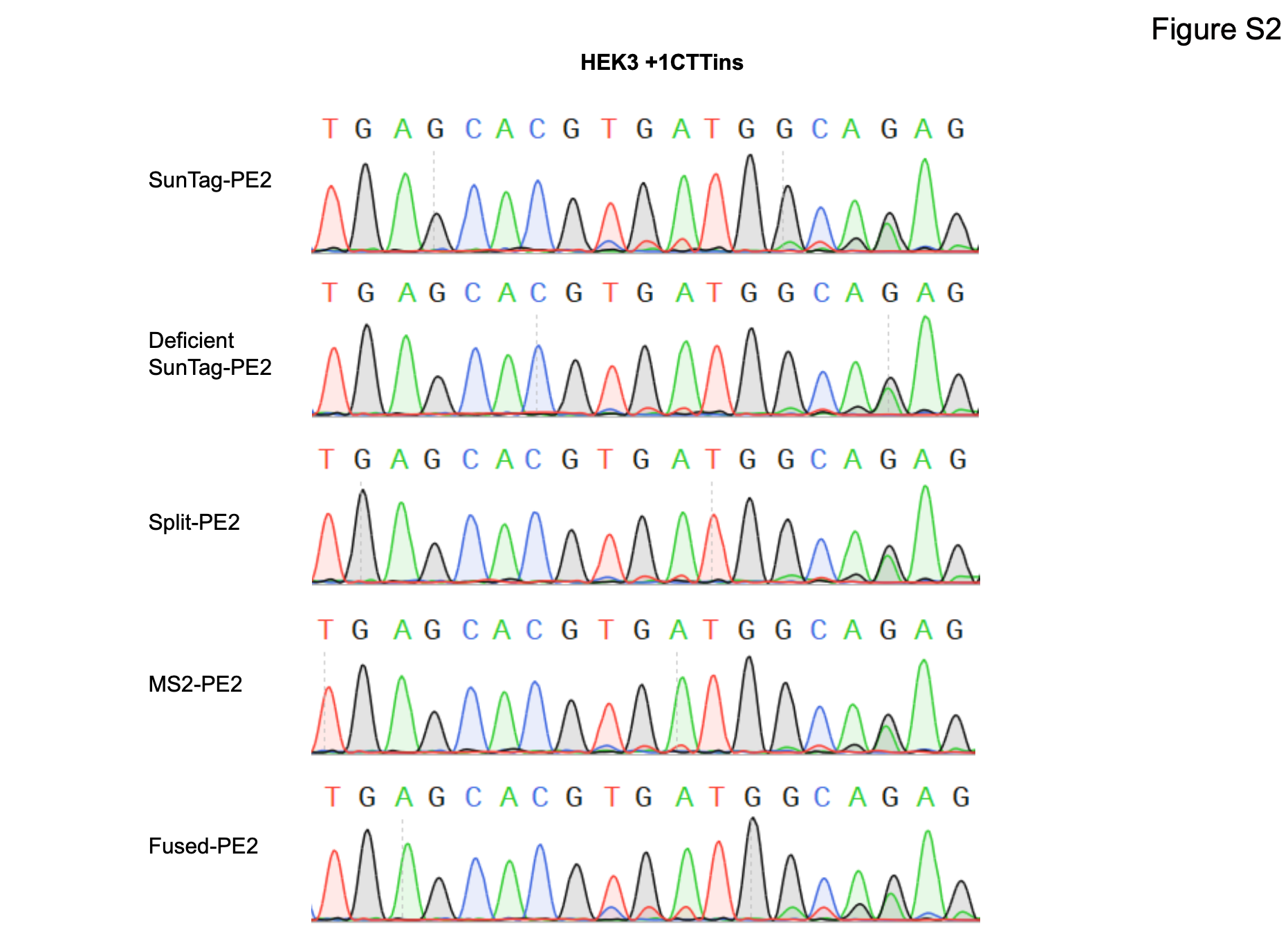

**Fig S2. Sanger sequencing results of HEK3 +1CTT insertion in Suntag-PE2, deficient SunTag-PE2, split-PE2, MS2-PE2, and canonical fused-PE2.**

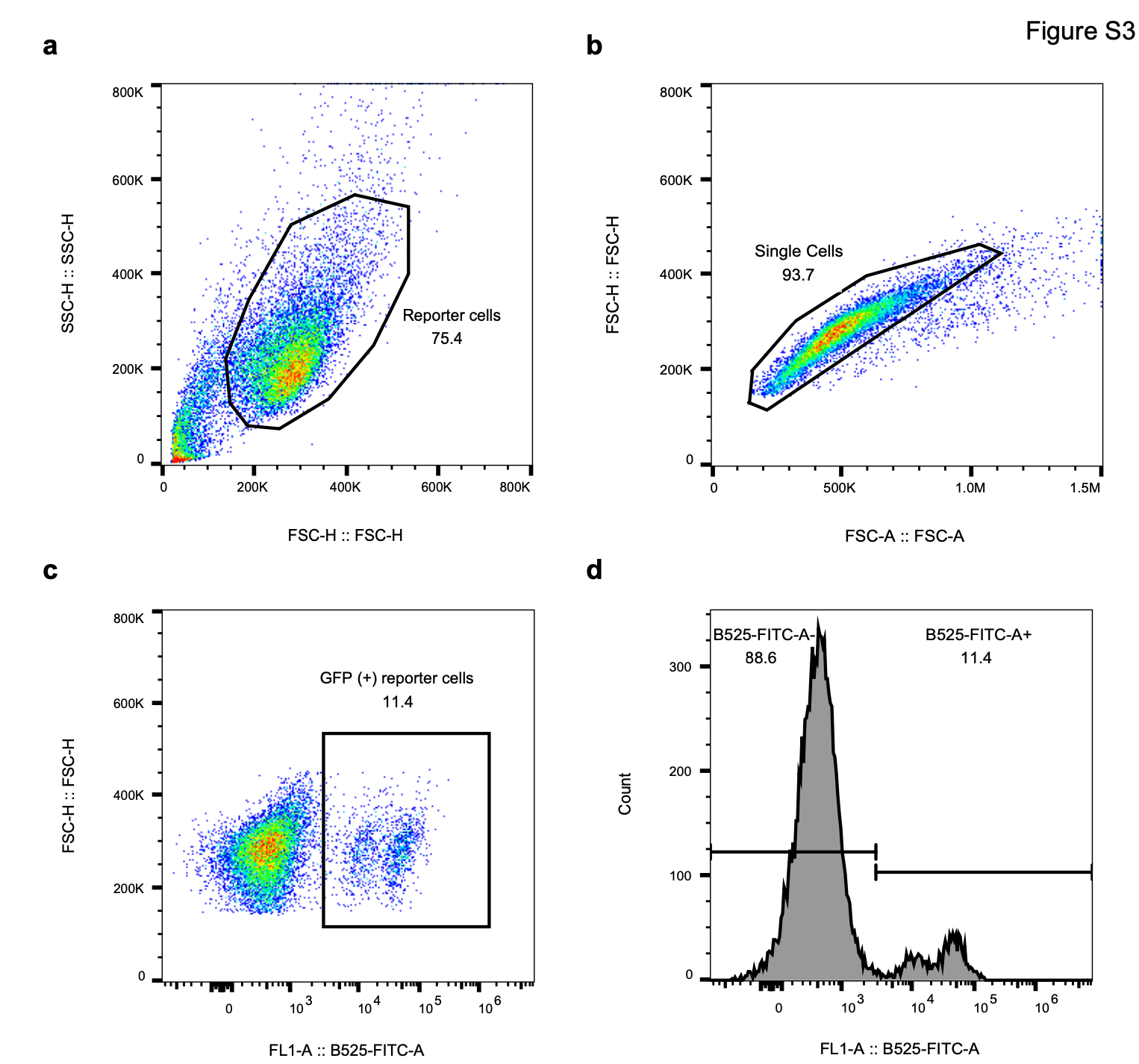

**Fig S3. FCM gating examples for GFPm reporter cells. a.** Gating for live cells. **b.** Gating for single cells. **c.** Gating for GFP positive cells. **d.** Histogram for GFP positive cells.

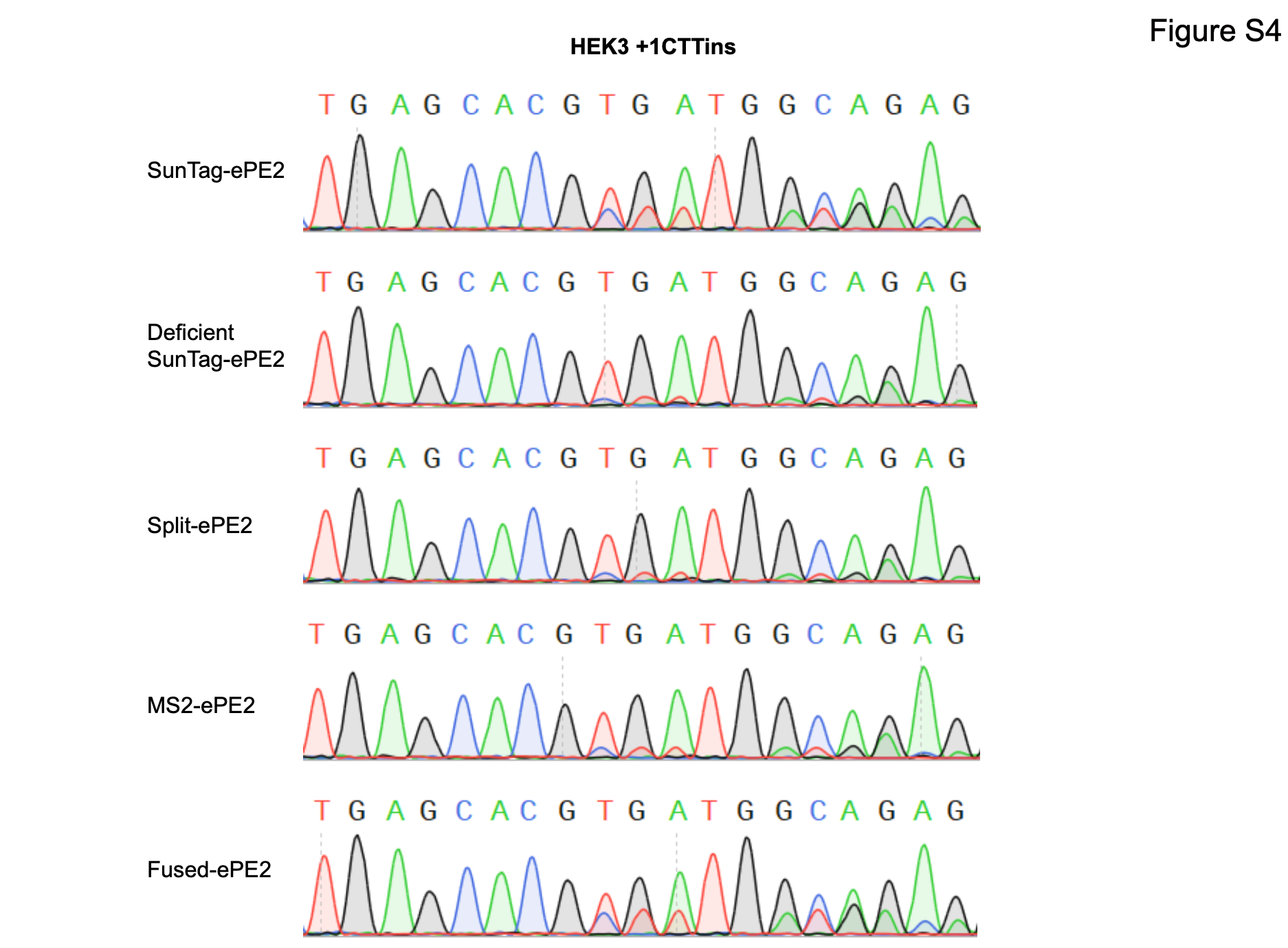

**Fig S4. Sanger sequencing results of HEK3 +1CTT insertion in Suntag-ePE2, deficient SunTag-ePE2, split-ePE2, MS2-ePE2, and canonical fused-ePE2.** The prime editing efficiency of different PE2 strategies in the ePE form was significantly higher than that in the PE form (Fig. S2)

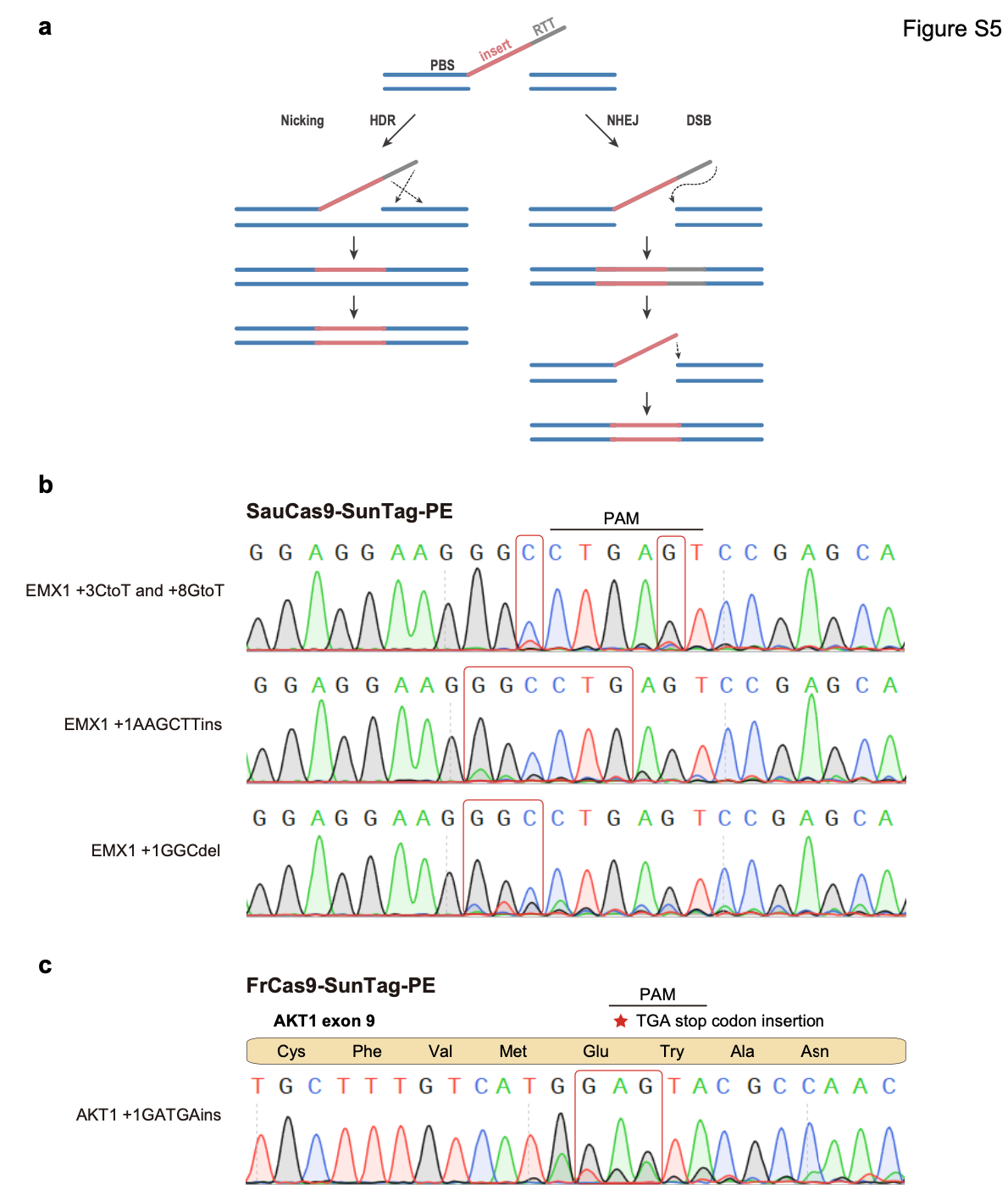

**Fig S5. DSB-SunTag-PE system of SauCas9 and FrCas9. a**, Schematic diagram of the repair difference between DSB-PE and nicking-PE. **b**, The sanger sequencing results of SauCas9-DSB-SunTag-PE in three types of desired mutations in EMX1 locus. **c**, The sanger sequencing results of FrCas9-DSB-SunTag-PE precisely inserted TGA stop codon in AKT1 exon 9.
