## Supplementary figures and images for "SunTag-PE: a modular prime editing system enables versatile and efficient genome editing"

### Extended Data Figure 1

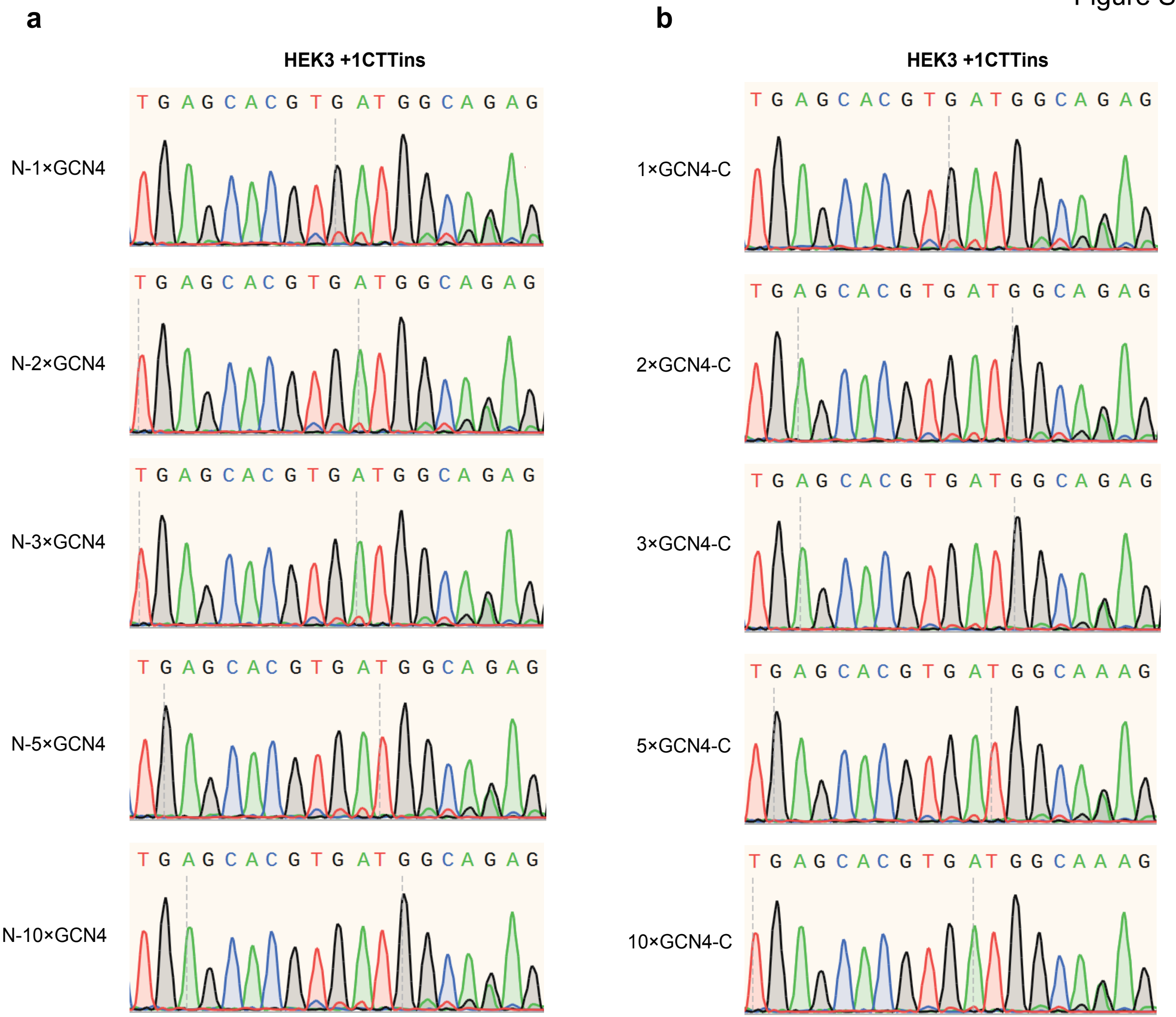

### Extended Data Figure 2

Figure S2

**HEK3 +1CTTins**

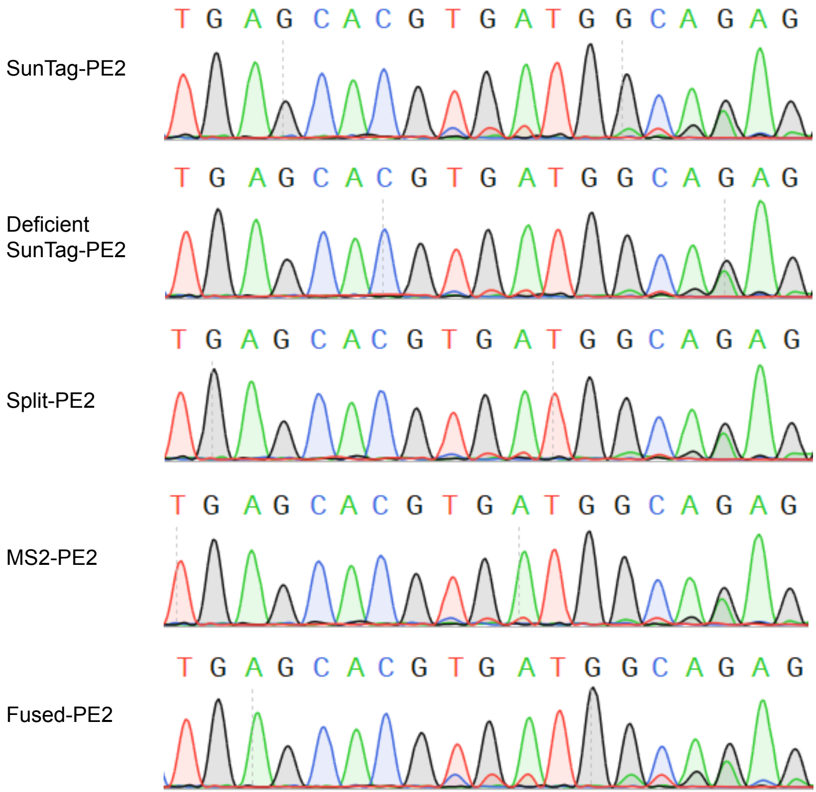

### Extended Data Figure 3

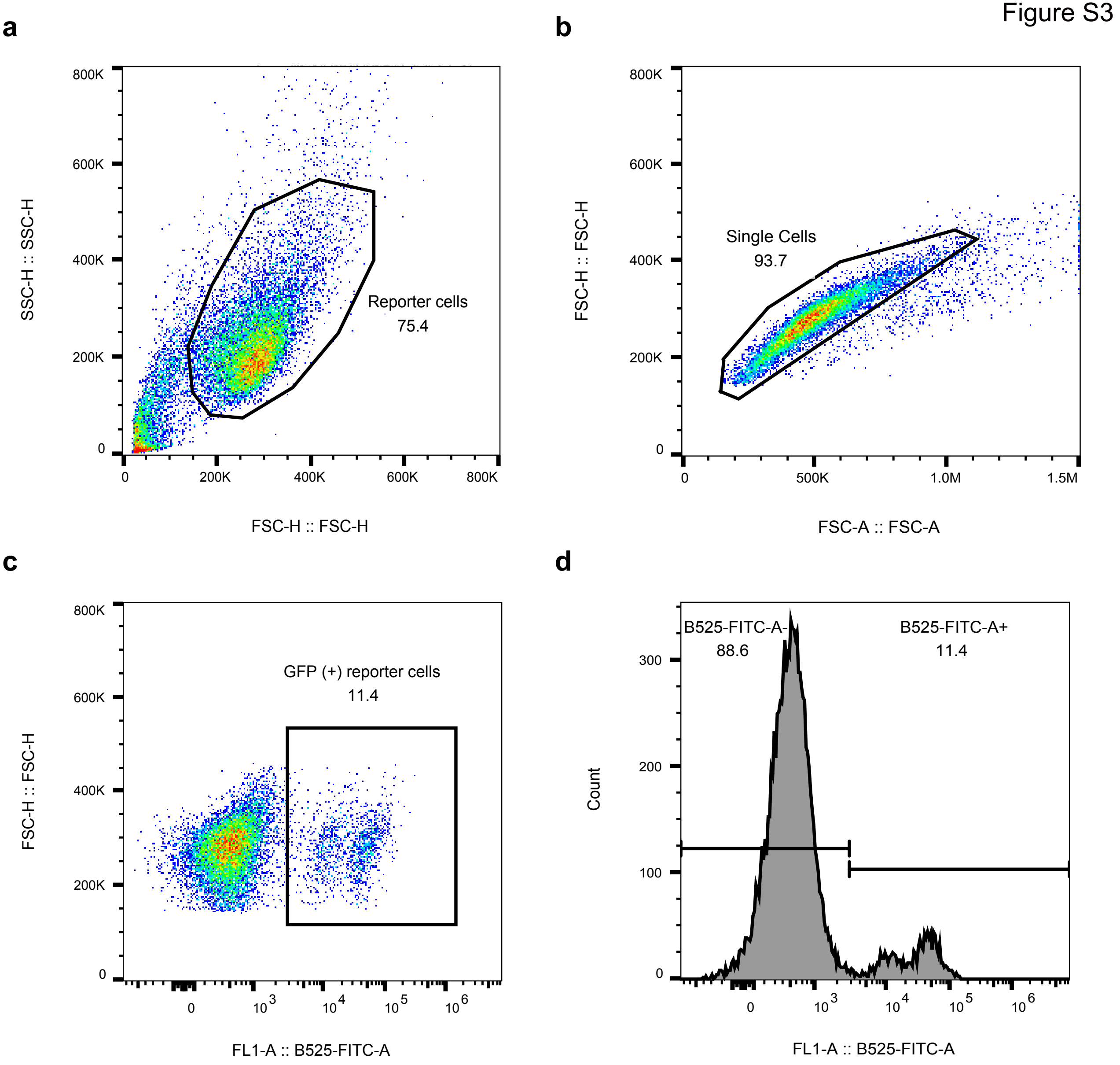

### Extended Data Figure 4

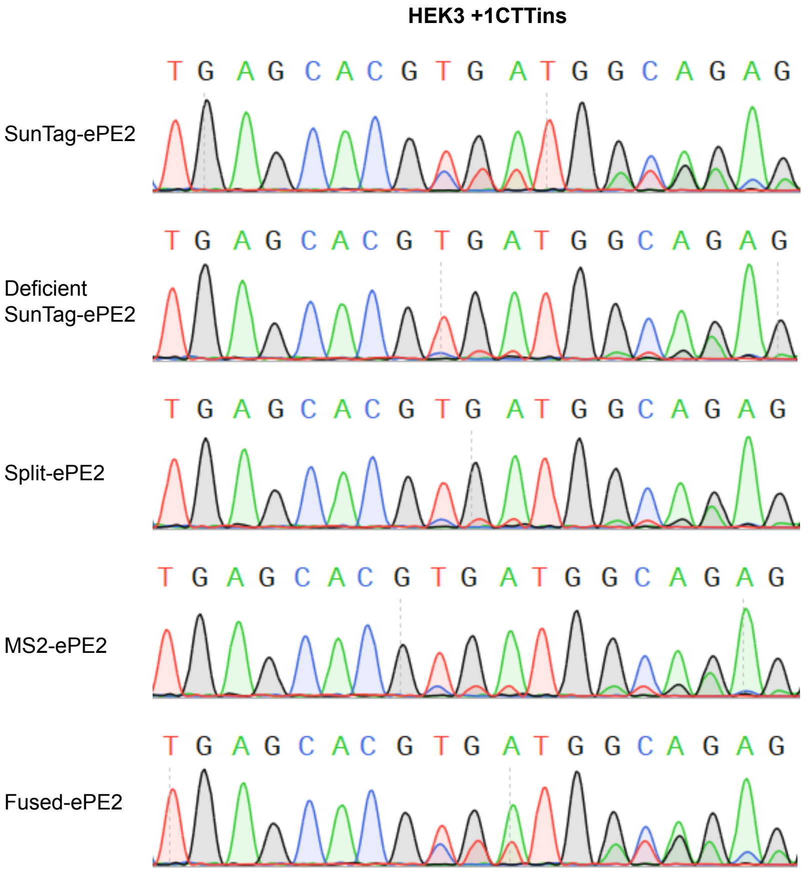

### Extended Data Figure 5

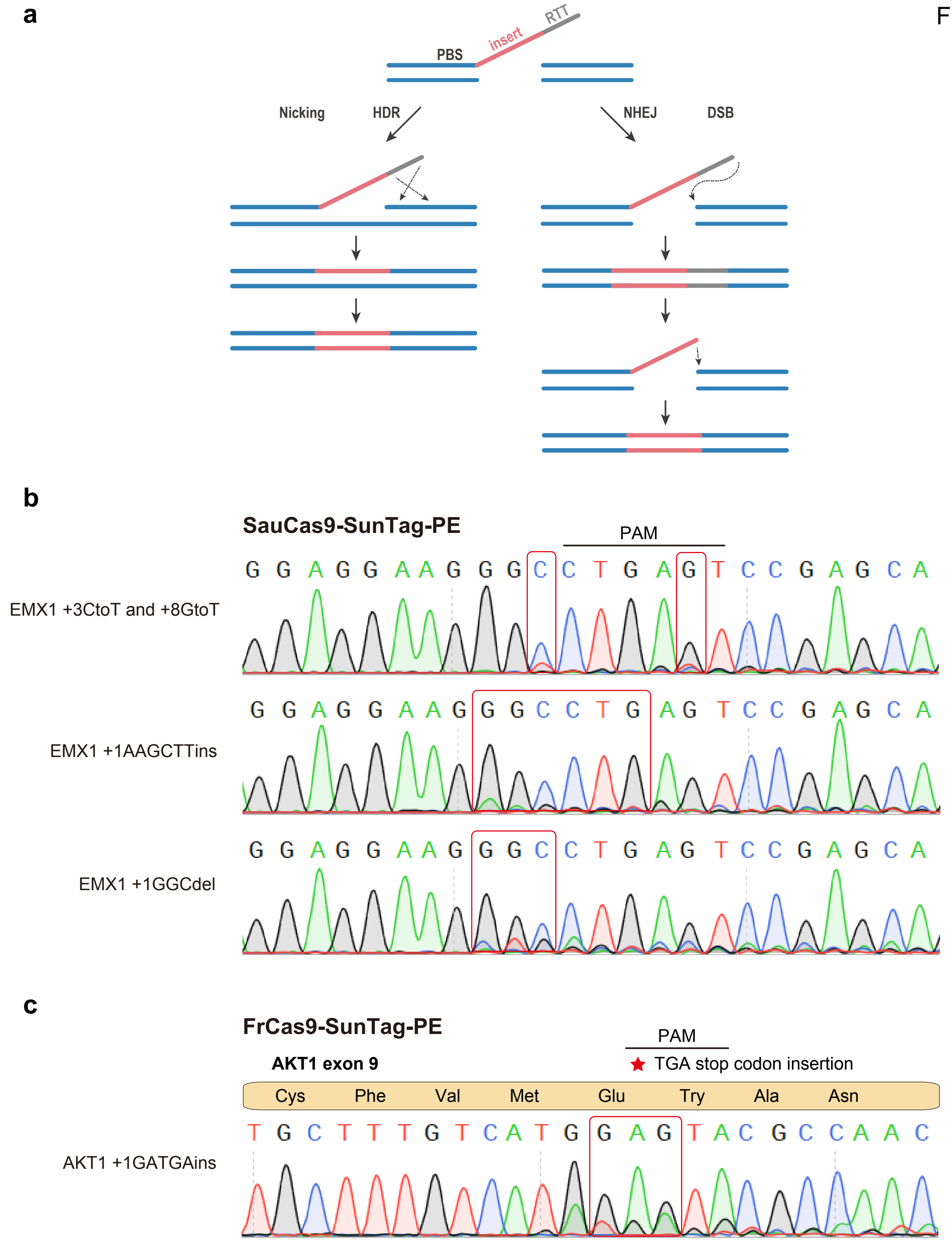
